## Supplementary Information for "Mechanism of DNA origami folding elucidated by mesoscopic simulations"

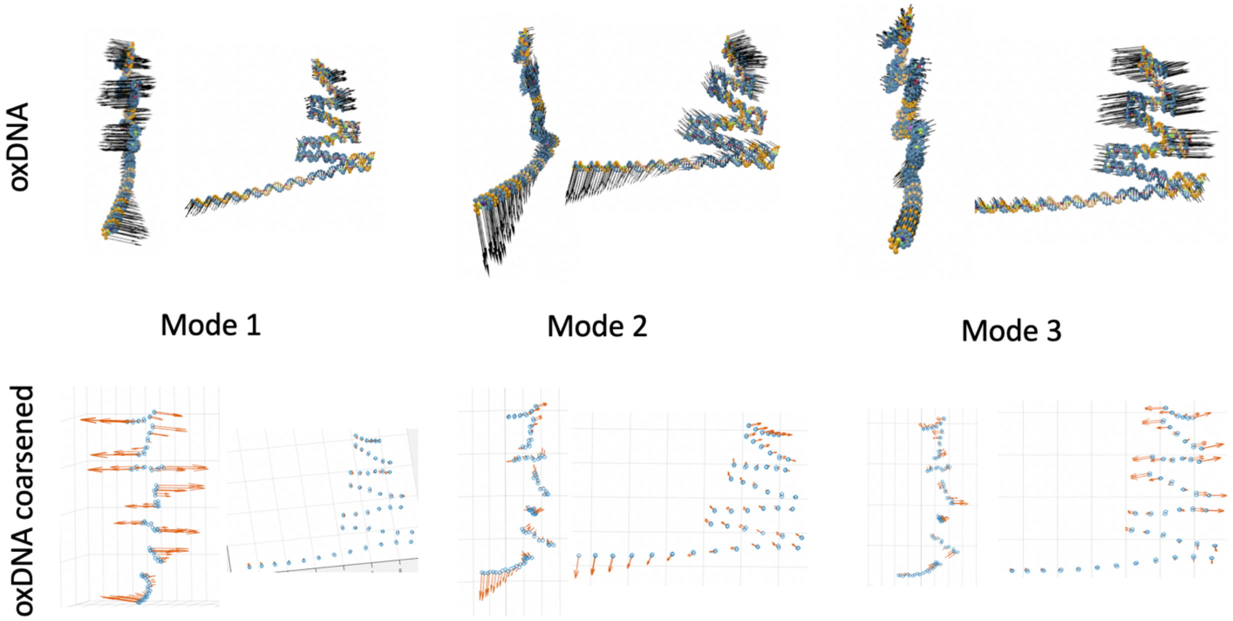


**Supplementary Figure 1:** First three principal components of sheet structure extracted from oxDNA simulations using single-nucleotide trajectory resolution compared to coarsened trajectory extracted from the same simulation. Results show good agreement, indicating that a model coarsened to 8 nucleotides per bead may be sufficient to capture the dynamics of DNA origami structures.

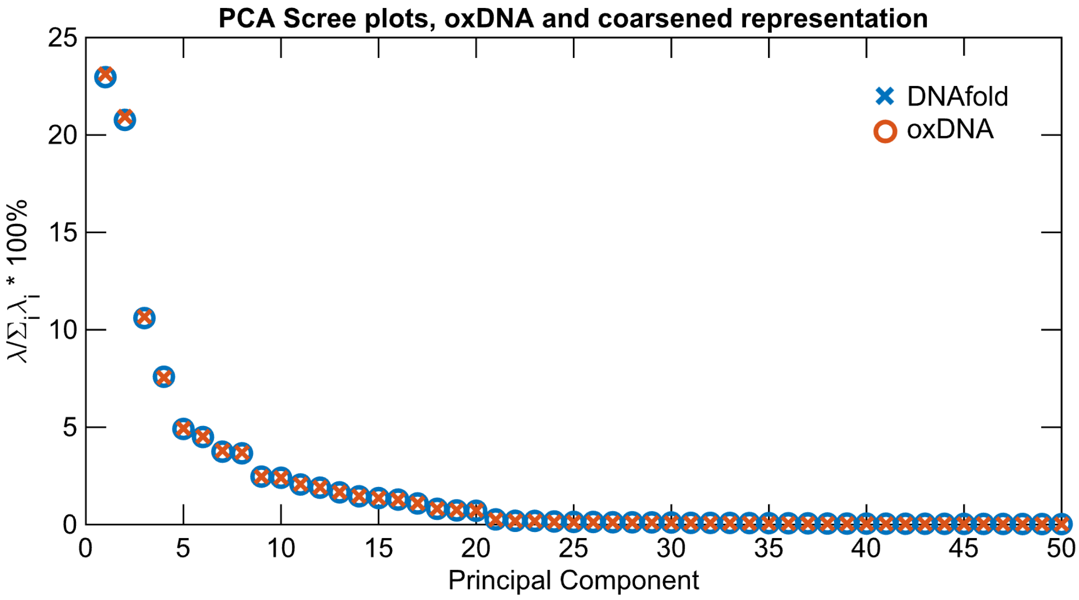


**Supplementary Figure 2:** Scree plots of first 50 principal components from PCA, oxDNA and mapped to the representation used in the DNAfold model.

**Supplementary Note 1: Viability of using Brownian dynamics for this model.**

The mass of each bead, which corresponds to 8 nucleotides, is approximately $2.64\,\mathrm{kDa}$ or $4.384 \times 10^{-24}\,\mathrm{kg}$. The Stokes friction coefficient of a spherical bead in a fluid is $\gamma = 6\pi\eta R$, where $R = 2.7\,\mathrm{nm}$ is the bead's radius and $\eta$ is the dynamic viscosity of the surrounding fluid (water), or approximately $2.545 \times 10^{-11}\,\mathrm{kg}/s$ at room temperature. It is typically considered appropriate to assume overdamped Langevin (Brownian) dynamics if the characteristic relaxation time is smaller than the timestep of motion, $m/\gamma < \Delta t_{\mathrm{step}}$. In our case, $m/\gamma \approx 1.723 \times 10^{-13}\,\mathrm{s}$ , which is approximately nine times smaller than our timestep $\Delta t_{\mathrm{step}} = 15 \times 10^{-12}\,\mathrm{s}$, and therefore it is acceptable to use Brownian dynamics for this model to more efficiently sample system dynamics.

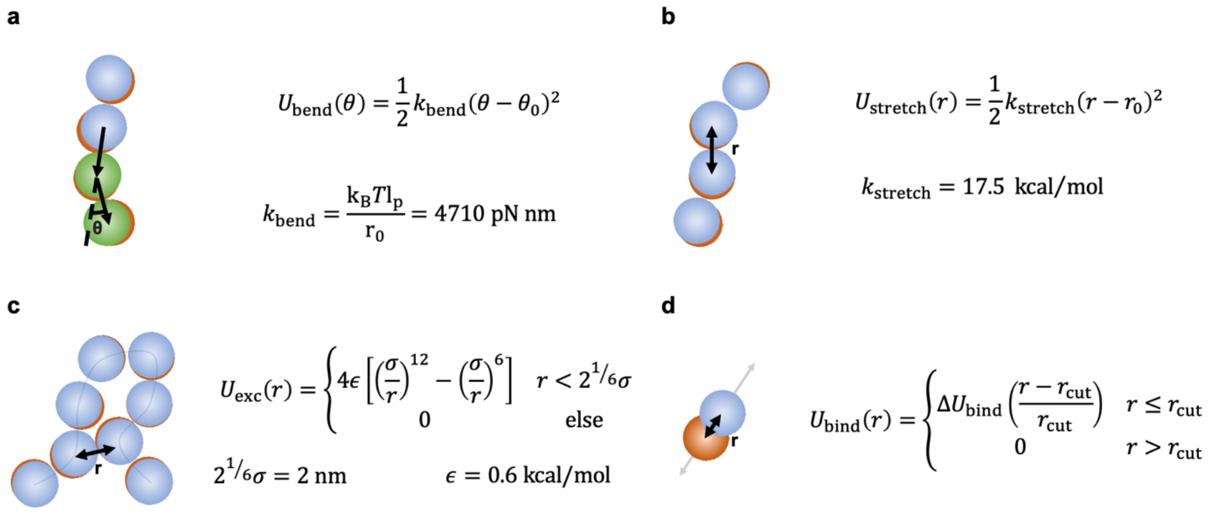


**Supplementary Figure 3:** Schematic of the four main potentials used for the DNAfold model, including (a) bending potential, (b) harmonic backbone potential, (c) WCA excluded volume potential, and (d) hybridization potential.

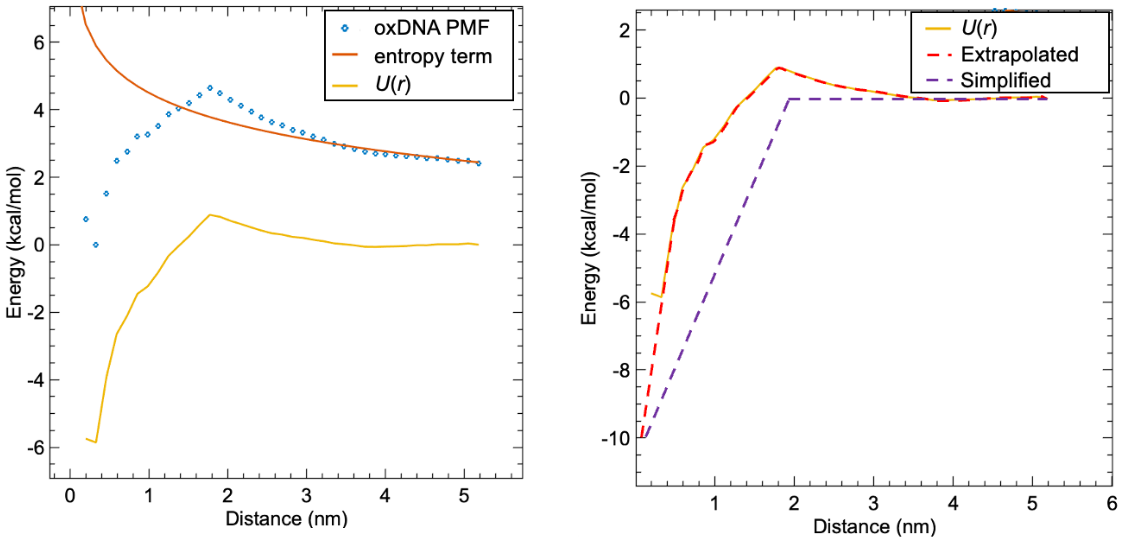


**Supplementary Figure 4:** The hybridization potential was derived using oxDNA-based umbrella sampling simulations of two complementary 8-nt sequences (using the sequence-independent model of oxDNA) being dehybridized. Distance denotes separation between centers of mass (COM) of the two 8-nt sequences and “Energy” denotes energy, PMF, or entropy. We then extracted the entropic term for separation, fitted it to the tail of the PMF curve, and subtracted this from the PMF to obtain hybridization energy as a function of COM separation.

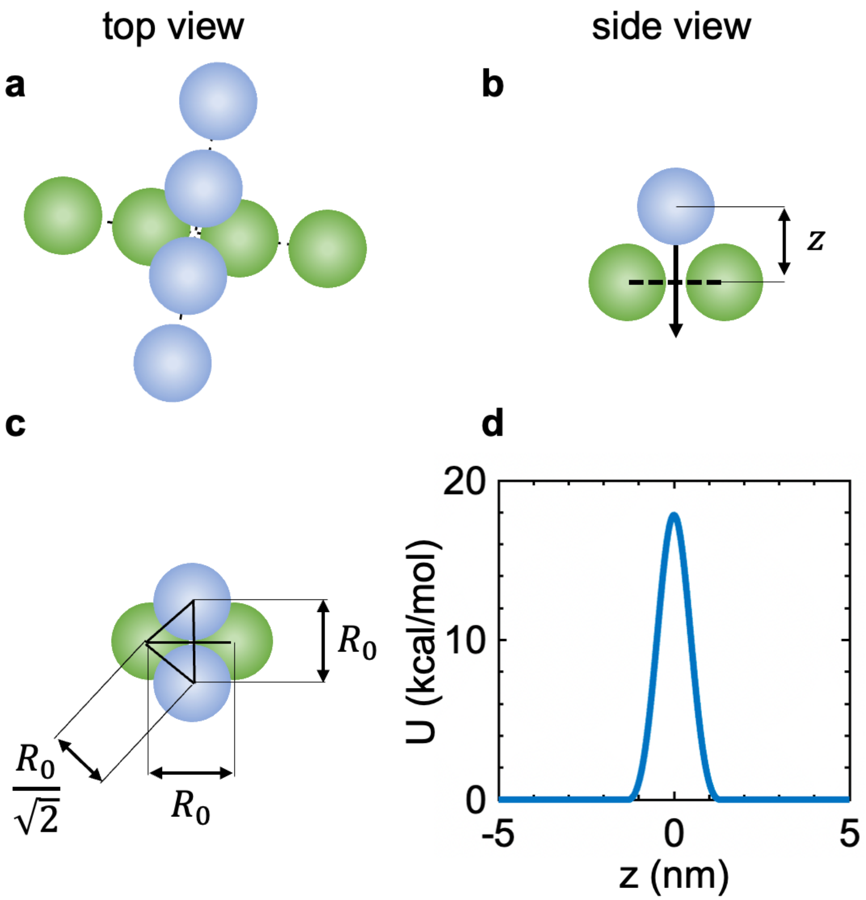


**Supplementary Figure 5:** (a-b) One primary function of the excluded volume potential is to prevent strand cross-through. The worst-case configuration for cross-through is one where a pair of consecutive particles approaches another pair of consecutive particles at a 90-degree angle forming a diamond pattern; this minimizes the excluded volume penalty to cross-through. (c) Geometry showing two beads located at $R_0/\sqrt{2}$ from two other beads (at height *z*=0) is the location of maximum potential and represents the minimum barrier to cross-through. (d) cross through potential arising from excluded volume interactions as a function of *z*, the height of one strand above the other, in this case depicted by blue above green.

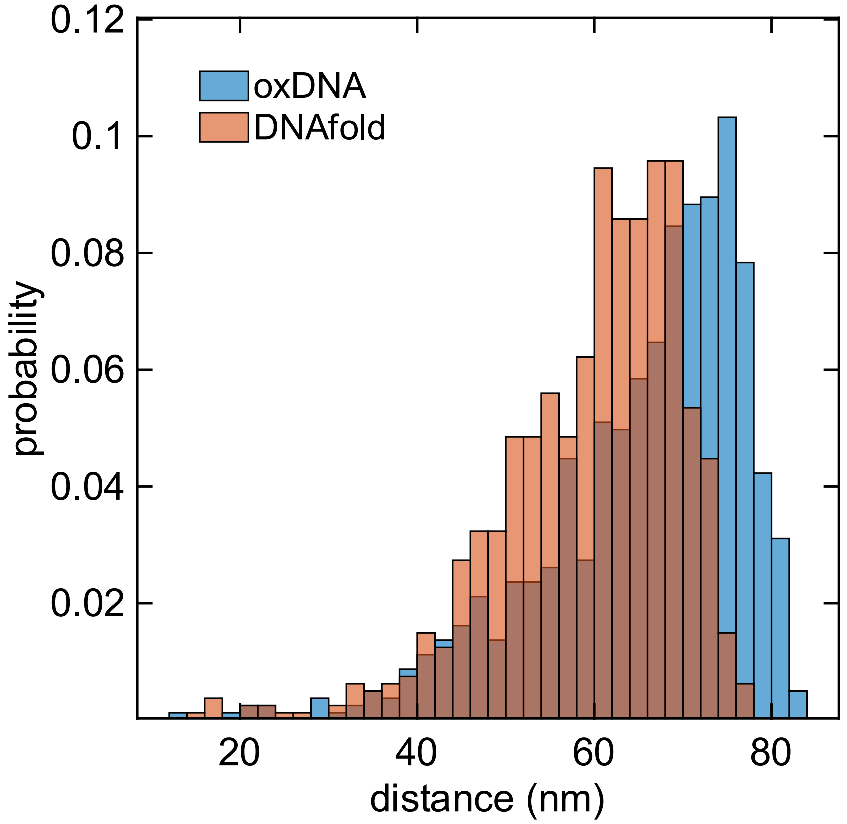


**Supplementary Figure 6:** End-to-end distance comparison between DNAfold and oxDNA for a 256-base pair section of dsDNA.

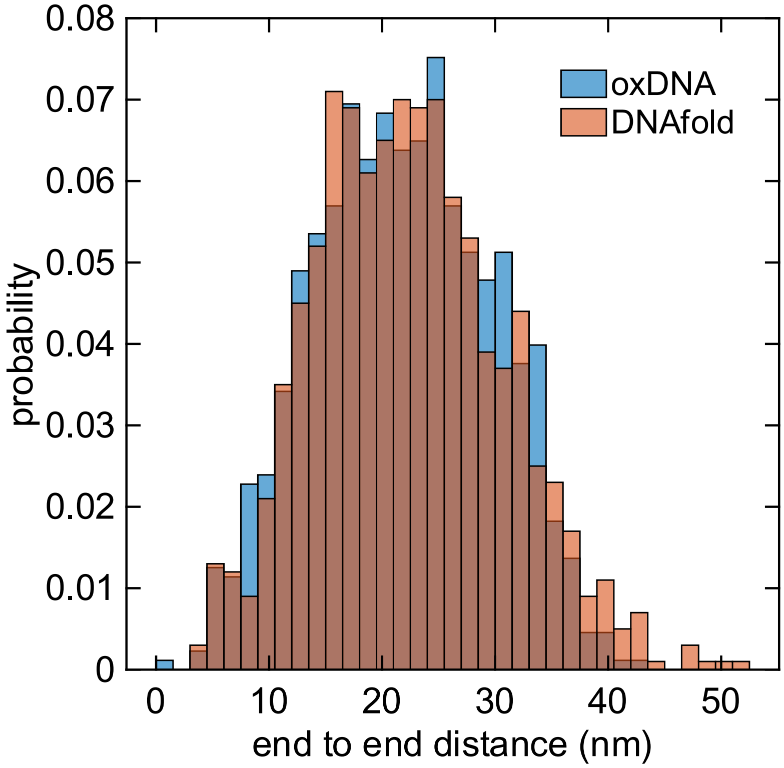


**Supplementary Figure 7:** End-to-end distance distribution comparison between DNAfold and oxDNA for a 256-nucleotide ssDNA section. Since this version of the model is sequence independent, we used polythymine as a reference.

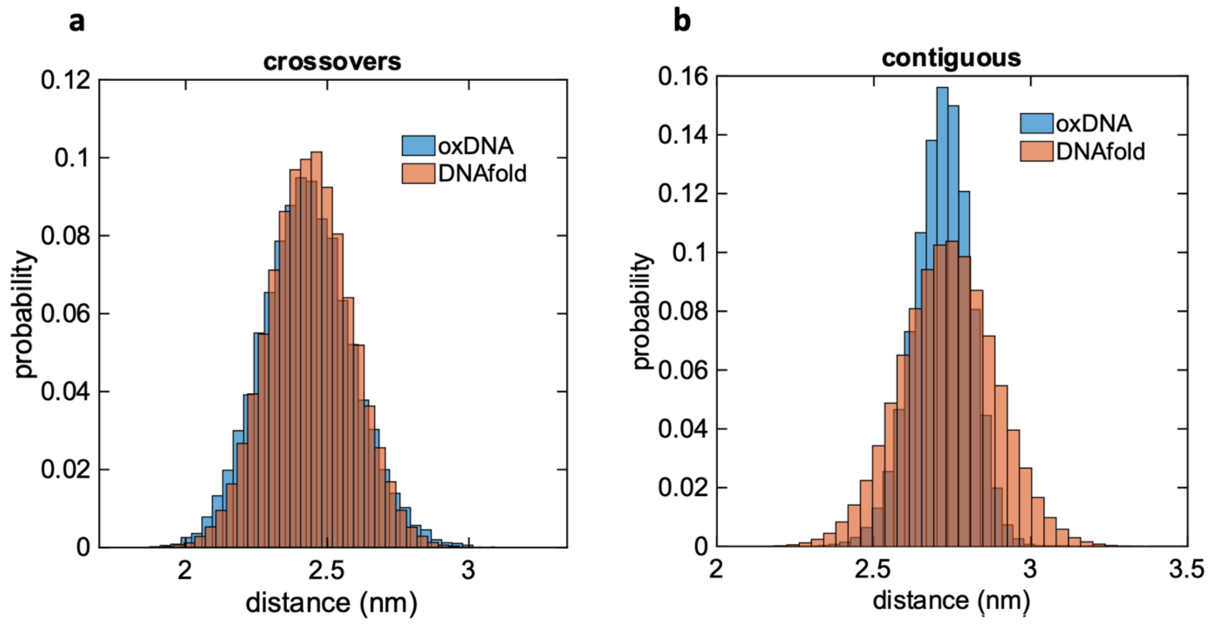


**Supplementary Figure 8:** Comparison of backbone length distributions from oxDNA to DNAfold for a fully assembled 768-nucleotide sheet structure. (a) distribution of separation distances between crossover beads. (b) distribution of separation distances between contiguous beads.

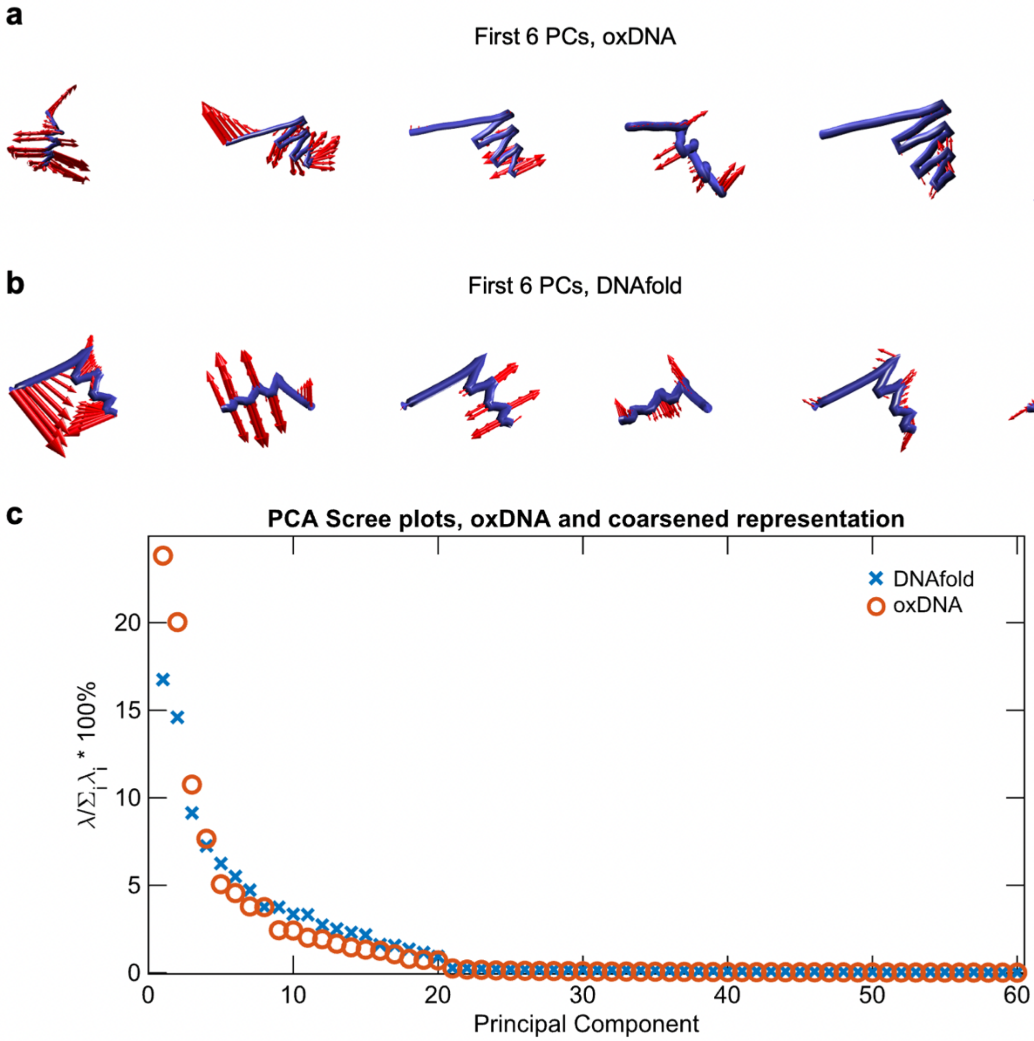


**Supplementary Figure 9:** Comparison of principal components and scree plots for oxDNA and DNAfold. (a) first 6 PCs for DNAfold sheet structure. (b) first 6 PCs for oxDNA sheet structure. (c) scree plots for both models mapped to same representation.

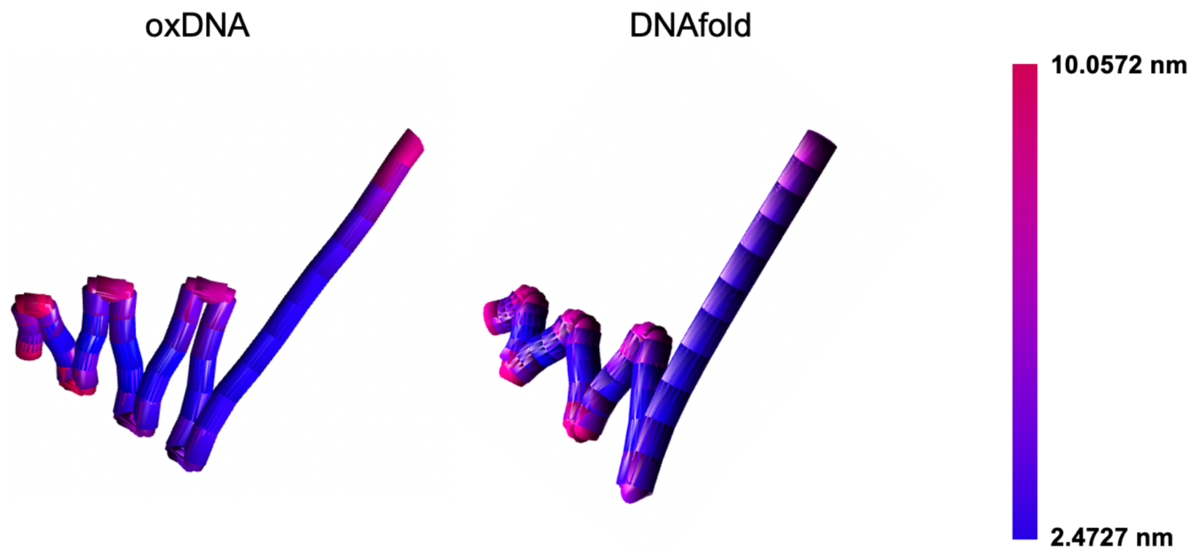


**Supplementary Figure 10:** RMSF comparison between oxDNA and DNAfold.

### Standard 4HB design

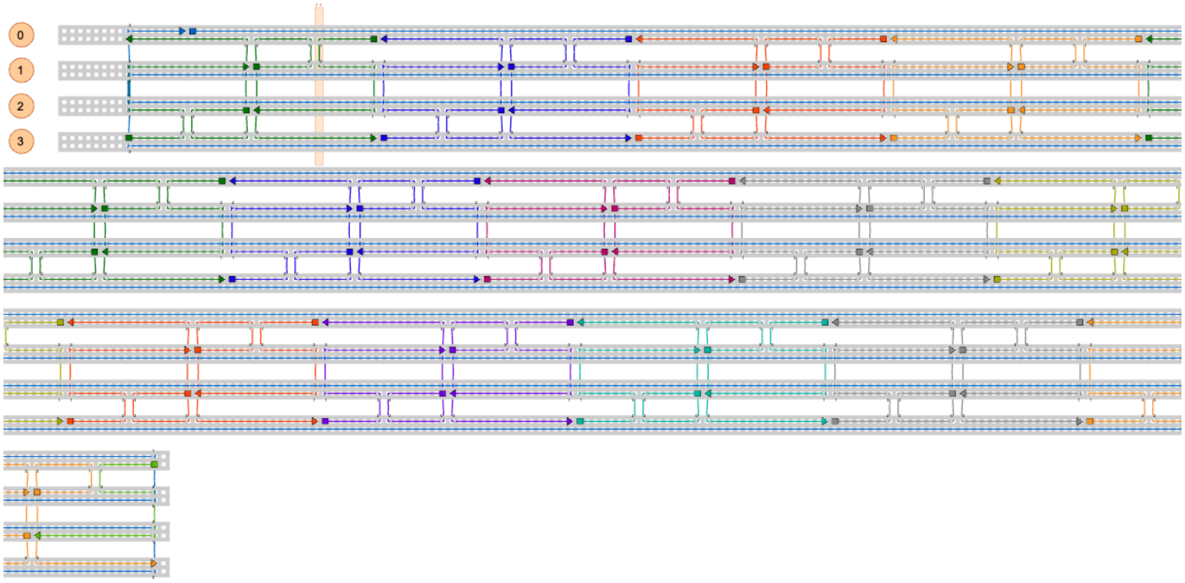


**Supplementary Figure 11:** caDNAno representation of standard four helix bundle (4HB) design.

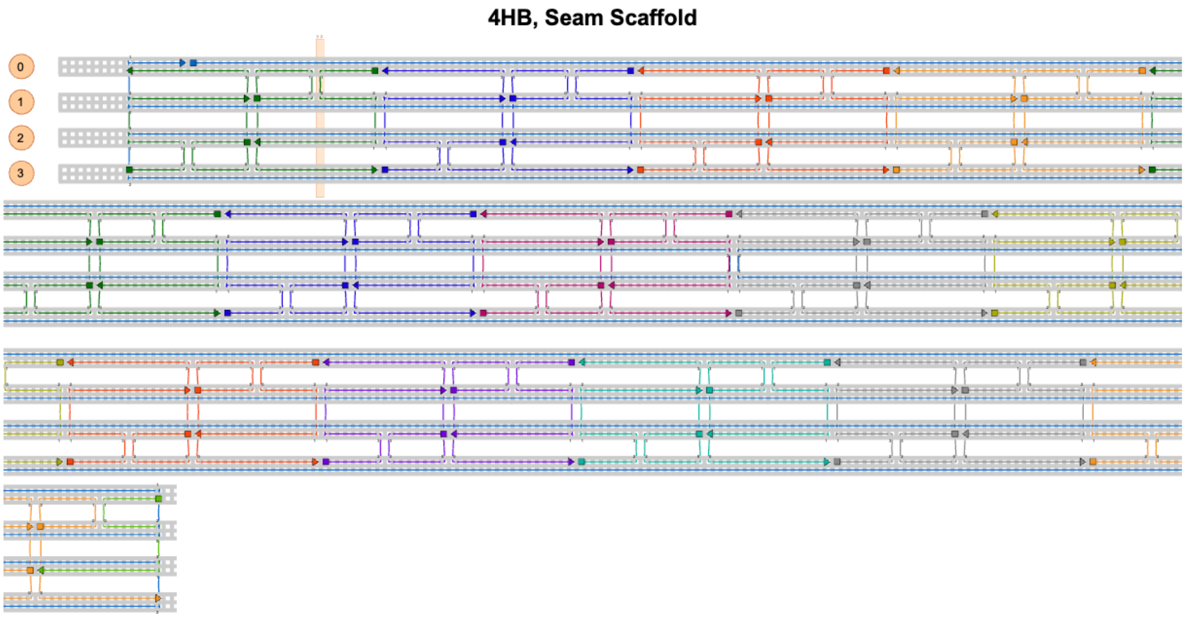


**Supplementary Figure 12:** caDNAno representation of 4HB design with seamed scaffold.

### 4HB, Winding Scaffold

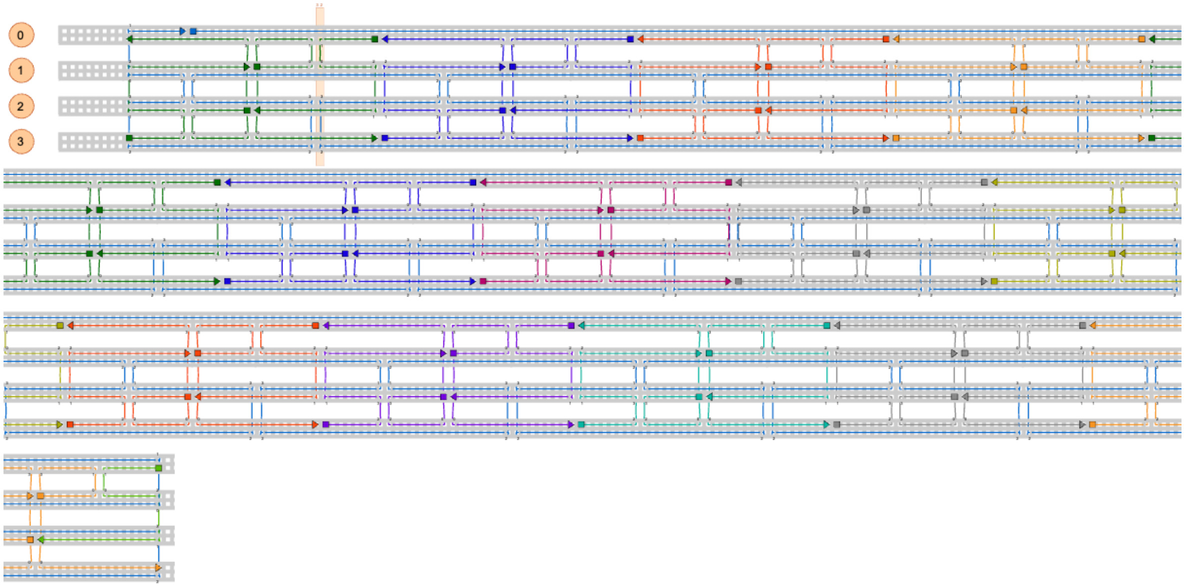


**Supplementary Figure 13:** caDNAno representation of 4HB design with winding scaffold.

### 4HB, Long Center Staples

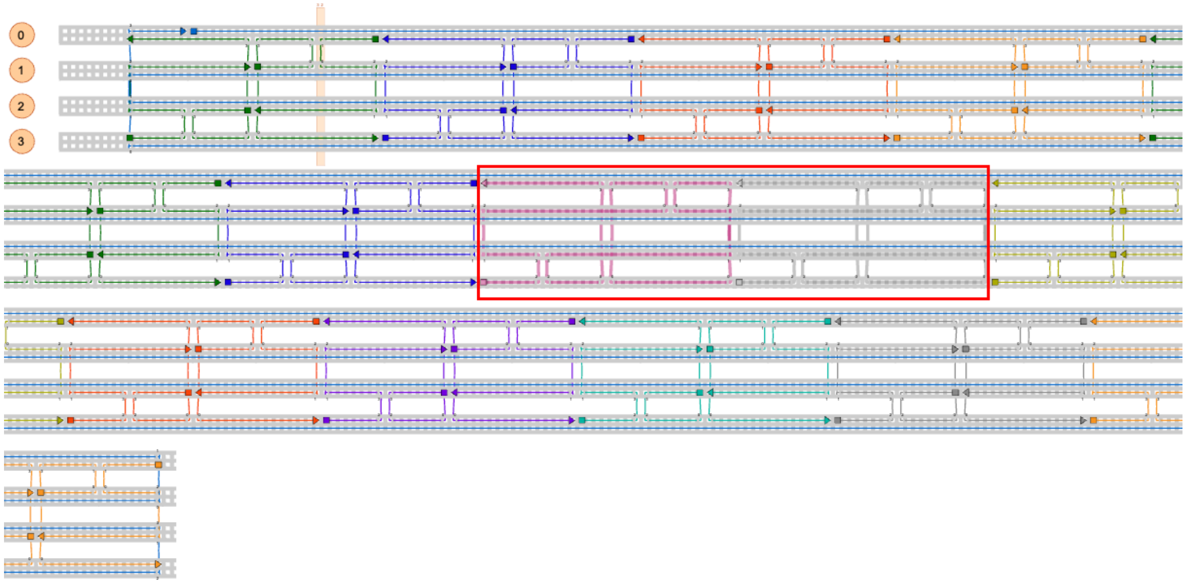


**Supplementary Figure 14:** 4HB with long center staples, shown in bold pink and grey and boxed in red.

#### 4HB, Long End Staples

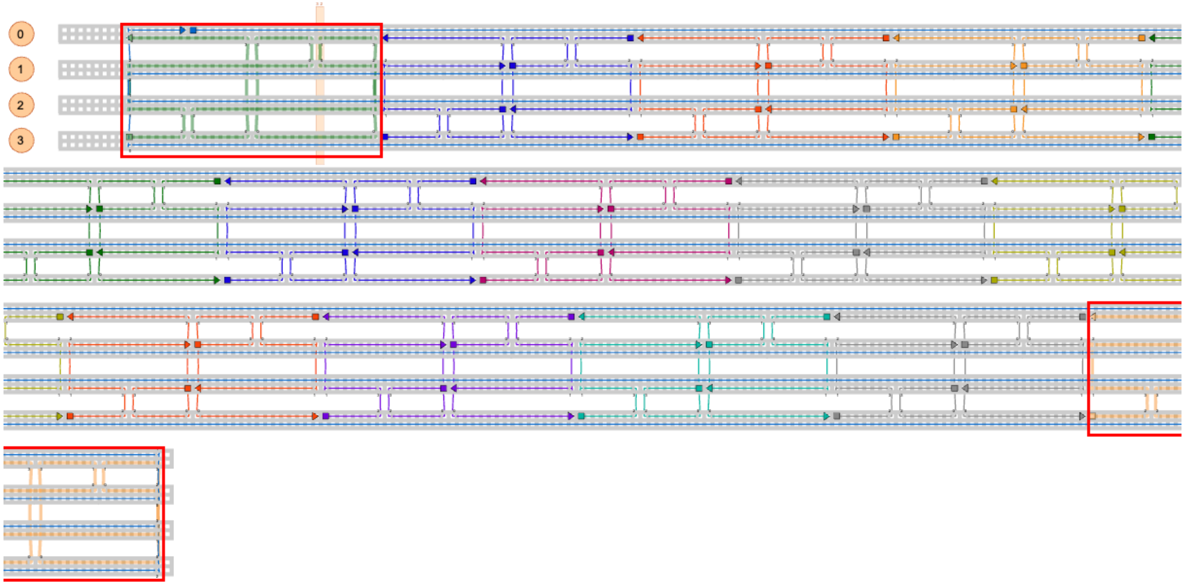


**Supplementary Figure 15:** 4HB with long end staples, shown in bold green and orange and boxed in red.

**Supplementary Videos 1-9: Self Assembly Animations for 4HB structures.**

Example folding animations can be found in the supplement with the following file names:

- *4HB_straight_routing_6kcal_mol.mp4*
- *4HB_straight_routing_8kcal_mol.mp4*
- *4HB_straight_routing_10kcal_mol.mp4*
- *4HB_straight_routing_12kcal_mol.mp4*
- *4HB_seam_routing_10kcal_mol.mp4*
- *4HB_winding_routing_10kcal_mol.mp4*
- *4HB_long_center_staples_10kcal_mol_collapse.mp4*
- *4HB_long_center_staples_10kcal_mol_slow.mp4*
- *4HB_long_end_staples_10kcal_mol.mp4*

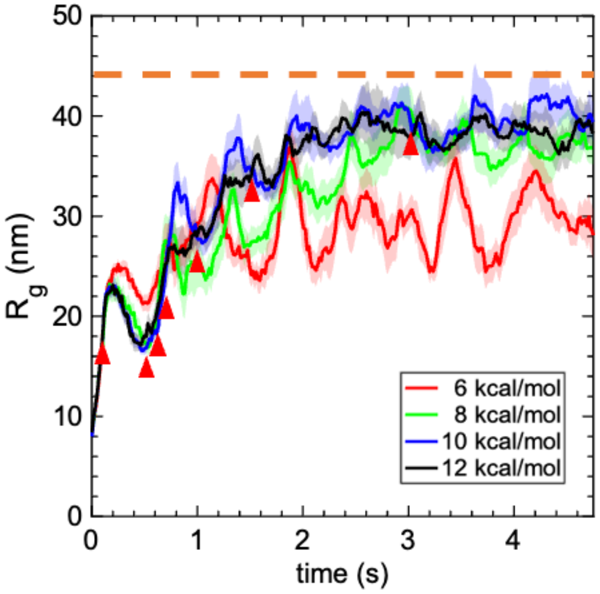


**Supplementary Figure 16:** Radius of gyration of 4HB structure during folding. The red arrowheads correspond to the snapshots from Figure 3 at 0.1, 0.5, 0.6, 0.7, 1, 1.5, and 3 seconds. The radius of gyration was calculated as $R_g = \sum_i \|(r_i - \bar{r})\|^2$, where $r_i$ is the Cartesian coordinate of particle $i$ and $\bar{r}$ is the centroid of the body of particles being analyzed. The orange dashed line represents the theoretical radius of gyration of a perfect 4HB in its idealized conformation based on the idealized caDNAno design. Fluctuations in $R_g$ are attributable to configurational fluctuations of the 4HB structure.

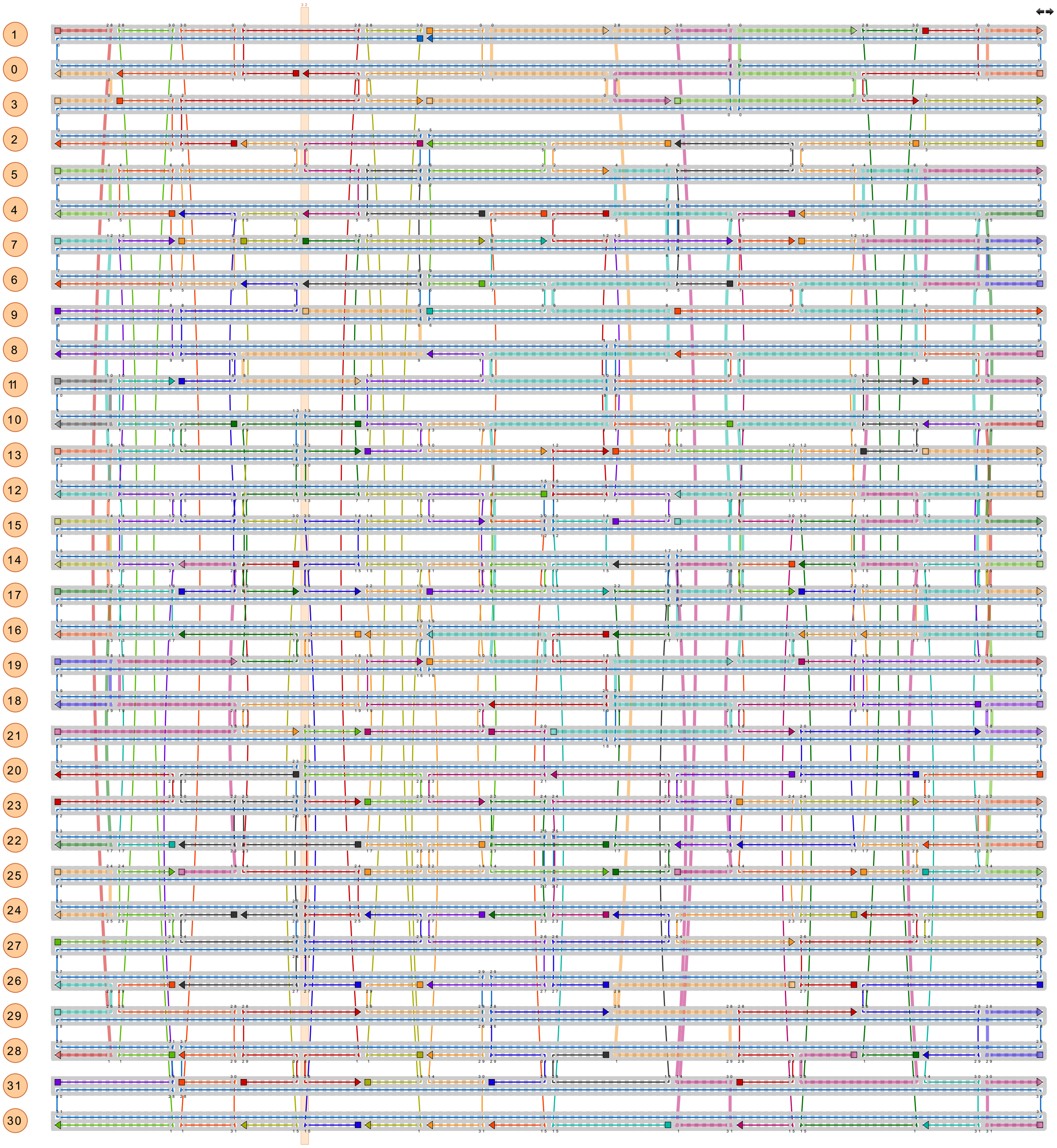

**Supplementary Figure 17:** caDNAno representation of 32 Helix Bundle design.

**Supplementary Videos 10-11: Self-assembly of 32HB.**

Folding animations can be found in the supplement with the following file names:

- *32HB_defect.mp4*
- *32HB_well_folded_with_staples.mp4*

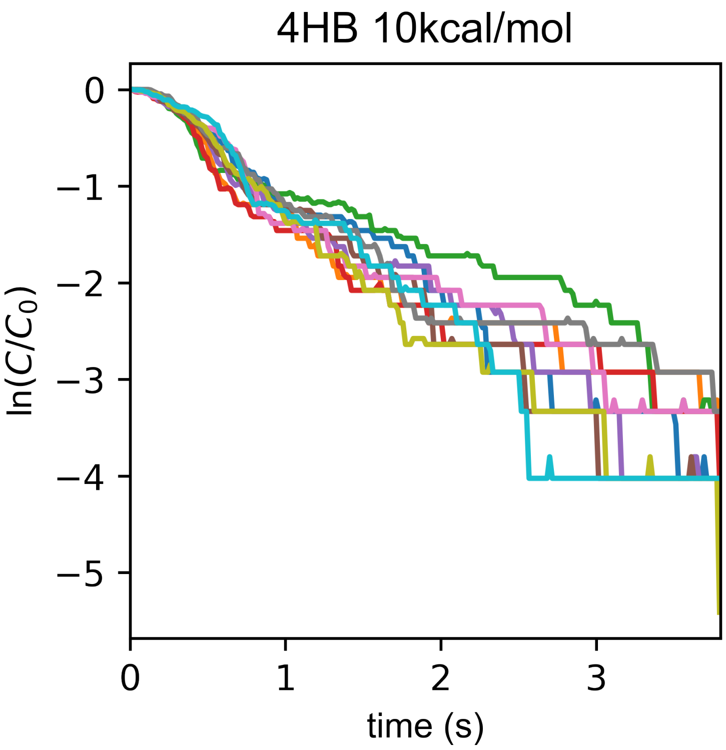


**Supplementary Figure 18**: Individual kinetic traces (ten in number) for 4HB with straight scaffold routing and hybridization strength of 10 kcal/mol per bead.
